# Oxytocin receptor antagonism during song tutoring in zebra finches reduces preference for and learning of the tutor’s song

**DOI:** 10.1101/2021.06.16.448133

**Authors:** Natalie R. Pilgeram, Nicole M. Baran, Aditya Bhise, Matthew T. Davis, Emily Kim, Sumin Lee, Carlos A. Rodriguez-Saltos, Donna L. Maney

## Abstract

In species with vocal learning, acquiring species-typical vocalizations relies on early social orienting. In zebra finches (*Taeniopygia guttata)*, for example, learning song requires dynamic social interactions with a “tutor” during an early sensitive period. The oxytocin system plays a central role in social orienting across species, yet it is unknown whether this system participates in the attentional and motivational processes that support vocal learning. Here, we tested whether blocking oxytocin receptors during exposure to tutors would impact learning from those tutors. Juvenile, song-naïve males were each tutored by two unfamiliar adults. During exposure to one tutor, juveniles were treated with oxytocin receptor antagonist (OTA) and during exposure to the other, saline (control). We found that OTA significantly reduced behaviors associated with approach and attention during tutoring sessions. Next, using an operant assay in which exposure to the two songs was balanced, we found that the juveniles preferred the control song over the OTA song. The developmental trajectory of preference for the control song resembled the pattern shown by father-reared birds choosing to hear their father’s song. Finally, the adult songs of the tutored birds more closely resembled control song than OTA song. The magnitude of this difference was significantly predicted by the early preference for the control song. Overall, oxytocin antagonism during exposure to a tutor seemed to bias juveniles against that tutor and his song. Our results suggest that oxytocin receptors play a role in socially-guided vocal learning in zebra finches, perhaps by affecting attention and motivation during tutoring.

---

Preferential orienting to social cues is critical for early social development and sets the stage for a lifetime of social learning. In humans, for example, the development of speech relies on early interactions with caregivers (Kuhl, 2007; Syal & Finlay, 2011). In songbirds, learning to sing is similarly social. Decades of experimental work has demonstrated that juveniles learn song most effectively from an adult “tutor” with which they can interact (reviewed by Gobes et al., 2019). In zebra finches (*Taeniopygia guttata*), nestlings separated from their father before fledging later produce songs with a number of aberrant features (Price, 1979). Juveniles that are able to hear adults singing, but not able to visually interact with them, do not learn their songs (Eales, 1987). Even before they begin to sing, juvenile male zebra finches form strong preferences for the song of their adult male caregiver (Rodriguez-Saltos et al., 2021; Fujii et al., 2021), and the magnitude of this preference predicts how well that song will be learned (Rodriguez-Saltos et al., 2021). These studies represent only a small fraction of a large literature suggesting that the mechanisms underlying song learning in zebra finches may share features in common with human vocal development—namely that song learning relies on social reward.

Oxytocin is well-known to mediate social reward. Although it is sometimes assumed to facilitate early orienting to a caregiver, including in humans, few studies have actually tested this hypothesis (Gordon et al., 2011; Hammock, 2014; Vaidyanathan & Hammock, 2017). In juvenile mice, administration of an oxytocin antagonist early in life reduced motivation to return to caregivers after a period of separation (Ross & Young, 2021). Similarly, in visually naïve chicks (*Gallus gallus*), administration of the avian homolog of oxytocin increased motivation to approach hens (Loveland et al., 2019). Thus, the role of oxytocin in attraction to social cues may be conserved across species and developmental stages. Accordingly, we and others have hypothesized that, in songbirds, oxytocin receptors (OTR) mediate attraction and attention to tutor song, facilitating song learning (Baran, 2017; Davis et al., 2019; Maney & Rodriguez-Saltos, 2016; Rodriguez-Saltos et al., 2021; Theofanopoulou et al., 2017).

Birds have a clear homolog of OTR (Gubrij et al. 2005; Leung et al., 2011); we will use the name “OTR” in keeping with established nomenclature (Kelly & Goodson, 2014; Theofanopoulou et al., 2021). OTR is expressed in many of the same brain regions in mammals and birds, including the lateral septum and auditory cortex (Leung et al., 2009; 2011); in songbirds it is also expressed in song control areas (Leung et al., 2009; 2011). We recently mapped OTR binding and mRNA in zebra finches throughout development, from day post-hatch (dph) five to adulthood, and found that this receptor is expressed in all of these regions throughout the entire period of song learning (Davis et al., 2019). In the song system, OTR is expressed at higher levels in males than in females (Davis et al., 2019). Together, these findings have formed the basis for our hypothesis that OTR mediates song learning by directing attention to tutors, facilitating the formation of social preferences, and promoting the development of selective encoding of tutor song (Baran, 2017; Davis et al., 2019; Maney & Rodriguez-Saltos, 2016; Rodriguez-Saltos et al., 2021).

To show that OTR activity contributes to the formation of social preferences during development, we administered an oxytocin antagonist to juvenile male zebra finches during early exposure to song tutors. Because zebra finches can produce accurate copies of songs after hearing as little as a few seconds per day over a few days (Tchernichovski et al., 1999; Deshpande et al., 2014), were able to limit tutor exposure precisely to times of pharmacological manipulation of OTR (London & Clayton, 2008). We hypothesized that pairing a song with OTR antagonism would reduce the reward value of hearing that song as well as the extent to which that song was ultimately learned.

## Results

### Oxytocin antagonism affected the pupils’ behavior during tutoring

Song-naïve juvenile male “pupils” (n=9) were tutored at ages 27-33 dph by two unfamiliar adult males in four separate sessions (Fig. 1), two sessions per tutor in alternating order. Pairs of tutors were assigned to each pupil according to the following criteria: (1) the two tutors sang at similar, high rates, (2) their songs were dissimilar to each other, and (3) their songs were also dissimilar to that of the pupil’s genetic father. Before each ∼1h tutoring session, pupils were treated either with oxytocin antagonist (OTA, see Methods) or saline (control) in counterbalanced order. Each treatment was always associated with the same tutor; in other words there was an “OTA tutor” and a “control tutor” (note that it was the pupils, not the tutors, that were treated). Here, we will use the terms “control song” and “OTA song” to refer to the songs heard by the pupil during each treatment condition. We balanced the number of times each pupil heard the two songs by playing recordings of a tutor’s song during the tutoring sessions if that tutor was not singing at the expected rate.

**Fig. 1.**
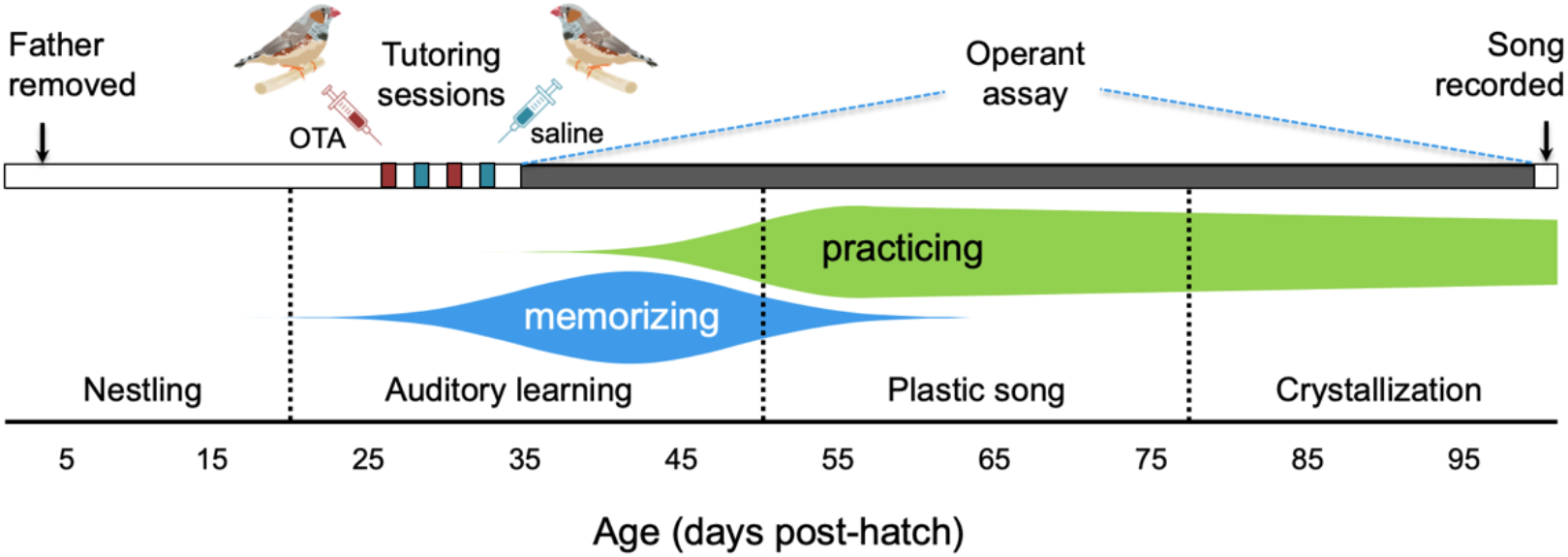
Experimental timeline and developmental trajectory of song learning in male zebra finches. Soon after fledging at ∼18 days post-hatch (dph), male zebra finches enter an auditory phase of song learning during which they memorize adult song (Eales, 1985) and start practicing singing (Zann, 1996). Singing rates increase dramatically around age 50 dph. At approximately the same time, juveniles enter the “plastic song” phase (Johnson et al., 2002; Kollmorgen et al., 2020), during which some memorization may still occur but ends by 65 dph (Eales, 1985). Song crystallization, during which the song takes its final form, begins by 77 dph and finishes by 90 dph (Johnson et al., 2002; Zann, 1996). In this study, male zebra finch “pupils” were reared in sound-attenuating chambers. Fathers were removed at four dph, such that pupils were thereafter isolated from male song until tutoring sessions commenced at 27 dph. During tutoring sessions, pupils were treated with either an oxytocin antagonist (OTA) during exposure to a tutor singing “OTA song” or treated with saline (control) during exposure to a tutor singing “control song” (see text). Treatments/tutors were alternated in counterbalanced order. At 35 dph, when they were nutritionally independent (Zann, 1996), the pupils were transferred to an operant chamber equipped with keys that played tutor songs according to a reinforcement schedule that allowed us to measure preference while balancing exposure to the two songs (Fig. 3; see also Rodriguez-Saltos et al., 2021). Song preference, namely preference for control song over OTA song, was measured daily while the pupils remained in the operant chamber, until 99 dph. We recorded the songs of the pupils at 101-103 dph and compared them to the control and OTA songs. Illustrations of finches by DataBase Center for Life Science, published under CC BY 4.0, https://creativecommons.org/licenses/by/4.0.

To test for effects of OTA on behavior during tutoring (Table 1), we quantified behaviors previously shown to predict song learning in this species (Chen et al., 2016; Houx et al., 2000). First, we scored approach to the tutor, operationalized in two ways. Time spent close to the tutor was defined as being within 12cm of the wall facing the tutor’s cage (designated as “the tutor zone”). Pupils tended to remain in the tutor zone regardless of treatment (Fig. 2A), so we were unable to detect an effect of OTA on proximity to the tutor. However, we did find that OTA significantly reduced the number of times the pupil approached the side of its cage closest to the tutor and touched its beak against that wall, which we scored as “pecks to tutor” (Fig. 2B). Thus, OTA did have an effect on observable interactions with the tutor.

**Table 1.** Effects of oxytocin antagonist (OTA) on behaviors during tutoring sessions. Behaviors were scored either as point counts per min or as a percentage of the trial, as indicated. The “tutor zone” was the area of the pupil’s cage closest to the tutor, defined as within 12 cm of the wall facing the tutor’s cage. “Pecks to tutor” is the number of times the pupil pecked at that wall.

|  |  | Control |  | OTA |  | Effect of OTA |  |  |
| --- | --- | --- | --- | --- | --- | --- | --- | --- |
| | Behavior | M | SD | M | SD | $\chi^2$ | $p$ | $d_{av}^\dagger$ |
| *Approach* | **Time in tutor zone** (percent of trial) | 69.4 | 28.7 | 79.9 | 27.3 | 1.68 | 0.195 | 0.42 |
|  | **Pecks to tutor/min** | 0.10 | 0.12 | 0.03 | 0.05 | 5.02 | **0.025\*** | **0.81\*** |
| *Attention* | **Flying** (two-footed wall contacts/min) | 0.22 | 0.62 | 0.19 | 0.46 | 0.02 | 0.878 | 0.11 |
|  | **Vocalizations/min** | 0.51 | 0.76 | 1.22 | 3.06 | 0.71 | 0.400 | 0.35 |
|  | **Preening** (percent of trial) | 2.96 | 2.54 | 1.61 | 1.51 | 9.64 | **0.002\*** | 0.64 |
| *Tutor* | **Song bouts/min** | 0.64 | 0.64 | 0.32 | 0.34 | 5.01 | **0.025\*** | 0.51 |

<sup>†</sup>Effect size is calculated as Cohen’s $d_{av}$ , which takes into account the within-subjects design (Cumming, 2012; Lakens et al., 2013).
\* $p < 0.05$ or $d_{av} > 0.8$ (large).

**Fig. 2.**
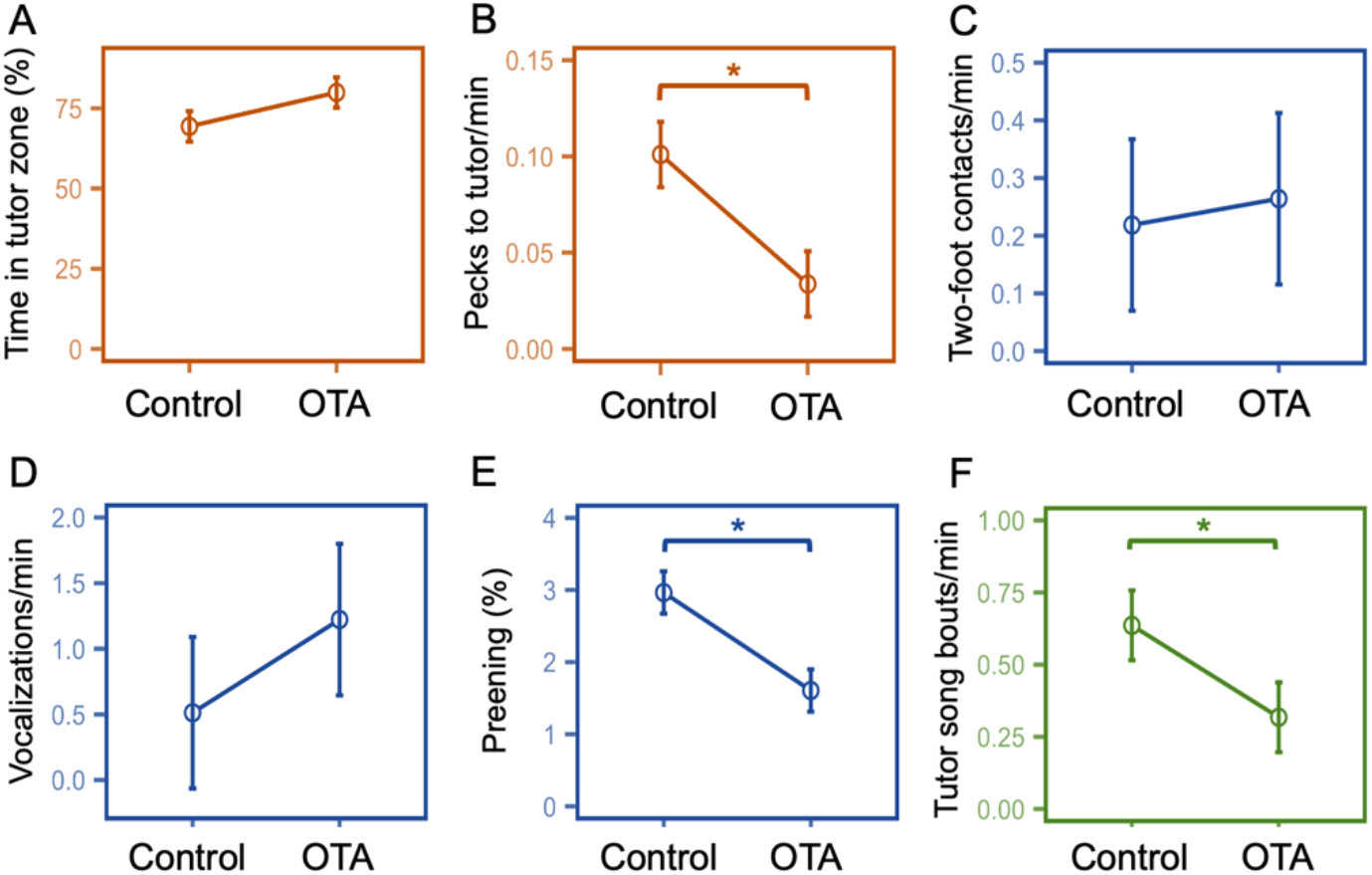
Effects of oxytocin antagonist (OTA) on behavior during tutoring. Approach-related pupil behaviors are shown in orange (A, B); attention-related behaviors are in blue (C-E); the graph in green (F) shows a tutor behavior. There was no effect of OTA on the time spent close to the tutor generally (A). Regardless of treatment, pupils tended to spend most of their time in a “tutor zone” within 12 cm of the side of their cage closest to the tutor. OTA significantly decreased the number of pecks to tutor (B), or the number of times the pupil touched its beak to the wall of its own cage closest to the tutor. There was no effect of OTA on flying, operationalized as two-foot contacts on cage walls (C), or on pupil vocalizations (D). OTA decreased the percentage of time pupils spent preening (E), which may be related to attention (Houx et al., 2000). Tutors sang at significantly lower rates when presented with OTA-treated pupils than when presented with control-treated pupils (F); however, tutor song bouts did not predict the pupils’ preferences or learning (see text). All panels show mean scores for saline and OTA trials. To accurately depict the within-subjects variation in the error bars, the between-subjects variation was removed using the *summarySEwithin* function in R (v4.04) which implements the method described by Cousineau (2005) and the correction factor described by Morey (2008). The plotted standard error of the mean (SEM) is therefore equal between conditions for each score. \**p* < 0.05. See Table 1 for statistics.

We also scored behaviors that may reflect attention to the tutor. Chen et al. (2016) found that song learning was enhanced when pupils sat quietly, not flying or vocalizing, during tutoring. Similarly, Houx et al. (2000) reported an inverse relationship between learning and both activity and vocalizations of the pupil. In this study, however, we detected no effect of OTA on flying, operationalized as the frequency of two-footed contacts with cage walls (Fig. 2C), or on vocalizations by the pupil (Fig 2D). We did, however, find an effect on preening, a behavior that occurs while birds are sitting quietly and that has been previously linked to the quality of song learning (Houx et al., 2000). OTA significantly reduced the amount of time the pupils spent preening (Fig. 2E), which may indicate less attentive behavior in the OTA condition.

### Treating pupils with oxytocin antagonist affected tutor song rate

OTA treatment of the pupils affected song rate in the tutors; the number of song bouts sung per minute by tutors when presented with OTA-treated pupils was significantly lower than when presented with control-treated pupils. We detected this effect two ways, first within-pupil (*X*^2^(1) = 4.355, *p* = 0.037, *d* _av_ = 0.71) and second, within-tutor (Fig. 2F; Table 1). Although we addressed any imbalances in tutor song bouts by playing recordings during tutoring, this result raised the possibility that effects of OTA on song preferences and learning could have been caused by hearing fewer *live* songs from the OTA tutors. We therefore checked for relationships between the number of live tutor songs and measures of preference and learning; we did not detect any such effects (see below).

### Oxytocin antagonism decreased preference for tutor song

After two tutoring sessions with each of the two tutors (at 27, 29, 31 and 33 dph), pupils were moved (at 35 dph) into a cage containing two keys that triggered playbacks of tutor song when pressed (Fig. 3A). One of the keys was more likely to play control song, that is, the song of the tutor paired with saline treatment. The other key was more likely to play OTA song. Each day, the pupil could press the keys to trigger 30 playbacks of each song. Because of our novel reinforcement schedule (Fig. 3B; see Methods), exposure to each song was balanced each day. Each pupil heard 30 playbacks of control song and 30 of OTA song, then the keys would trigger no further playbacks until lights-on the following day. After being transferred to the operant cages, the pupils engaged with the task and were exhausting the daily quota of 60 playbacks (30 of each song) within 5.5 ± 4.7 days (M ± SD), a time frame similar to that reported by Rodriguez et al. (2021) for juveniles that were not naïve to song. Pupils remained in the operant chamber, completing the task each day, until 99 dph. One pupil was excluded from the study because of low levels of key-pressing; two others became ill shortly after being transferred to the operant cage and were also excluded, leaving six pupils that completed the key-pressing portion of the study.

**Fig. 3.**
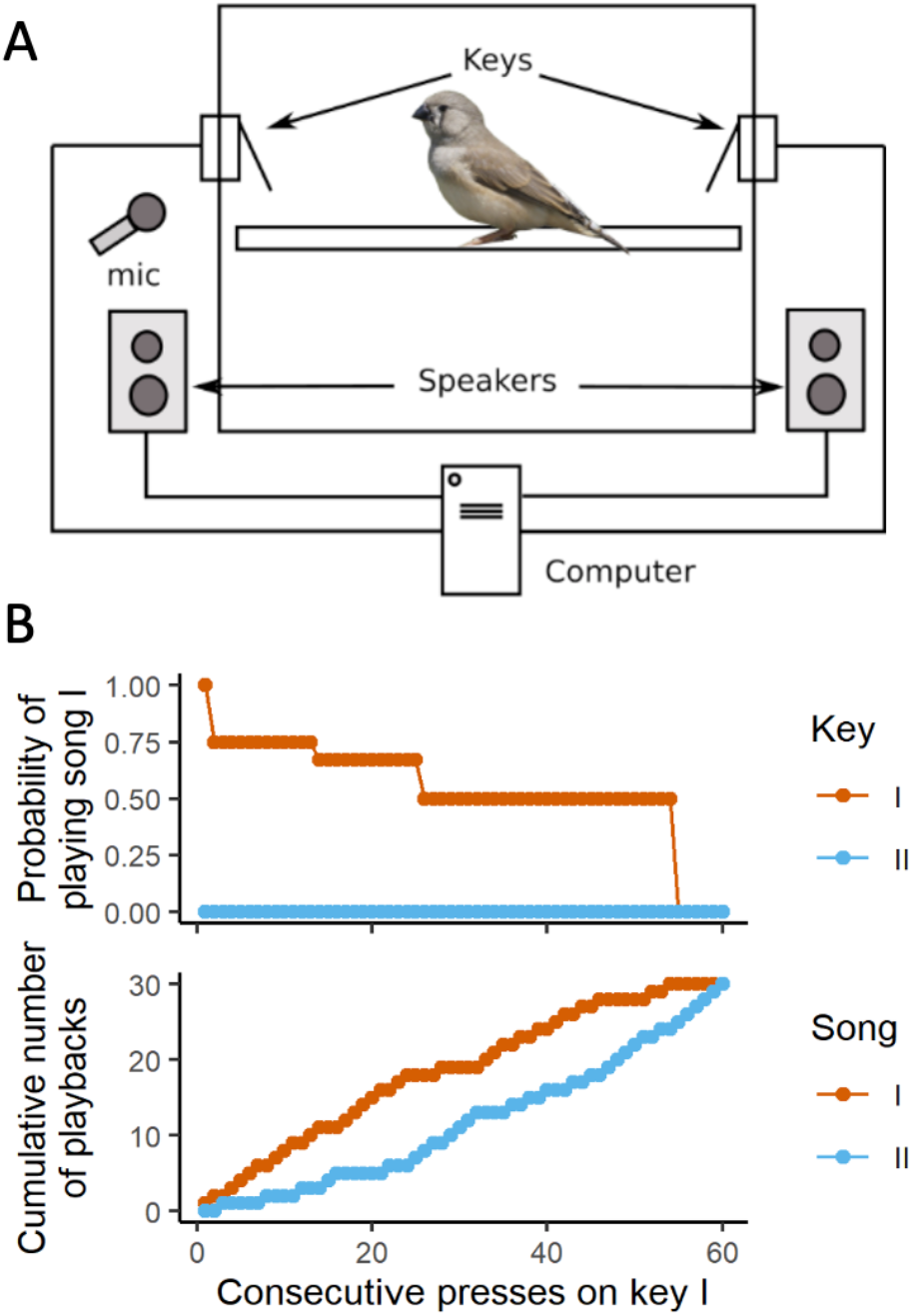
Key-pressing set-up and reinforcement schedule. (A) The operant chamber consisted of a 14×14×16 inch cage, inside which two keys were placed on opposite walls. One key was associated with playback of control song and the other with playback of OTA song. Outside the cage, one speaker assigned to each key played the songs. (B) demonstrates the reinforcement schedule. Keys I and II are associated with a higher probability of playing song I and song II, respectively (Rodriguez-Saltos et al., 2021). Here we present an extreme example to illustrate how the schedule is able to balance exposure despite a strong preference. In this scenario, the bird prefers song I and presses key I only. The probability of playing song I by pressing that key is high at the beginning of the session, to help the bird form the association between that key and the song. As the bird keeps pressing key I, the probability decreases step-wise from 1 to 0.5, to prevent song II from lagging far behind in the playback count. This decrease, however, is not enough to balance exposure, and therefore, if the bird switches keys, key II plays only song II until the playback count of song II is balanced with song I. After enough presses on key I, song I eventually reaches a quota of 30 playbacks and ceases to be played. Afterwards, only song II is played, until that song also reaches the quota. Importantly, there is always a large difference between the keys with respect to the probability of hearing the preferred song. When the key associated with preferred song is playing that song only 50% of the time, the other key plays non-preferred song 100% of the time. The code used in this study, as well as an updated version, is available from Rodriguez-Saltos et al. (2021). Photo of finch by Lip Kee Yap, shared under the Creative Commons Attribution-Share Alike 2.0 Generic license.

We estimated the strength of the preference for control song over OTA song daily by calculating the proportion of presses on the key associated with control song. A generalized additive model showed that preference changed over time (*p* = 1.41 e-5) (Fig. 4A). On average, control song was significantly preferred over OTA song (preference for control song > 0.5; 0.5 outside 95% confidence interval [CI]) between ages 40 and 47 dph, during the auditory phase of learning. At ∼51 dph, average preference shifted toward OTA song, reaching an initial, significant peak at 55-59 dph, during the onset of plastic song. Preference for OTA song was significant again from 82-95 dph, during song crystallization. This pattern, with presses for control song reaching an early peak followed by a peak in presses for OTA song, was seen in all of the individual trajectories (Fig. S1). The pattern closely resembled that exhibited by normally-reared males, which on average prefer to hear their father’s song during auditory learning but then come to prefer a different song later, during plastic song and crystallization (Rodriguez-Saltos et al., 2021).

**Fig. 4.**
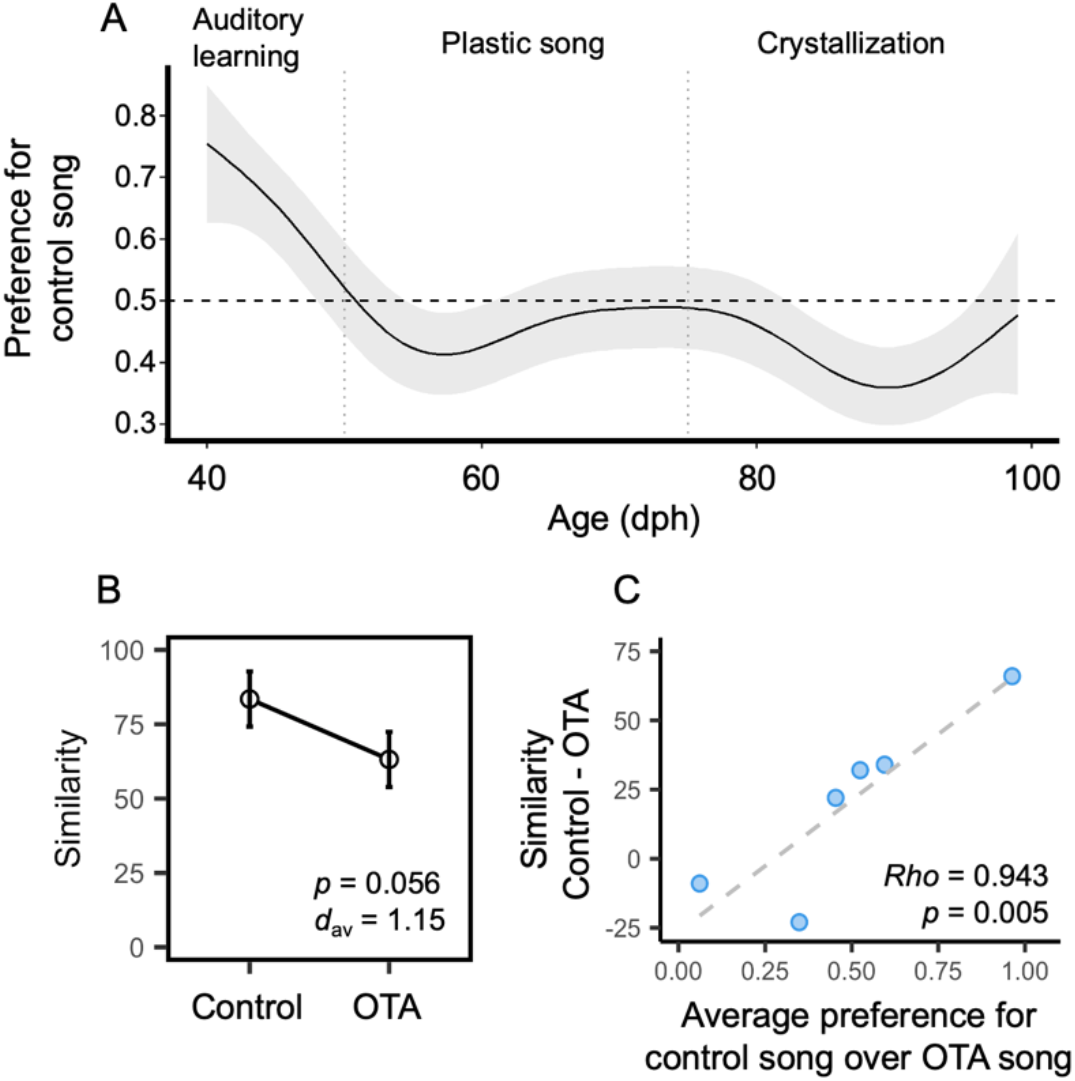
Effects of oxytocin receptor antagonist (OTA) during tutoring on song preference and learning. (A) Generalized additive model showing the trajectory of song preferences over the entire period of vocal development. The plot shows the average trajectory of the preference for the song of the tutor in the control condition, or ‘control song’. The horizontal dashed line at 0.5 indicates chance; values above the line indicate preference for control song and values below indicate preference for OTA song. The shaded area indicates the 95% confidence interval (CI). Vertical dashed lines indicate boundaries between developmental phases (see Fig. 1). On average, preference for control song was significant during the auditory learning phase (CI does not include 0.5). (B) Average similarity between pupil song and tutor song was higher in the control condition than the OTA condition. The effect size (Cohen’s *d*_av_) was large, at 1.15. (C) The preference for control song over OTA song (plotted on the X-axis) significantly predicted the degree to which birds learned control song better than OTA song (Y-axis).

The number of live songs sung by the control tutor during the tutoring sessions did not significantly predict average preference for control song, either over the entire trajectory of vocal development (*Rho* = 0.543; *p* = 0.266) or during the auditory learning phase (*Rho* = 0.429; *p* = 0.397). Thus, we do not have evidence that the early preference for control song was caused by hearing more live song from control tutors.

### Oxytocin antagonism inhibited song learning

The pupils’ crystallized songs were recorded at 101-103 dph for comparison with both the control song and OTA song. Similarity scores, calculated with Sound Analysis Pro 2011 (Tchernichovski, 2011), were higher for control song than for OTA song (Control: M = 83.5 ± 11.726 SD, OTA: M = 63.167 ± 23.617 SD) (*X*^2^(1) = 3.662, *p* = 0.056, *d*_av_ = 1.15; Fig. 4B). Note that although the p value was slightly above 0.05, the effect size was large. The number of live tutor songs heard during the tutoring sessions did not significantly predict learning (Control *Rho* = 0.029; *p* = 0.957; OTA *Rho* = -.486; *p* = 0.329; Fig. S2). The latter result was consistent with the findings of Houx and ten Cate (1998), who noted no correlations between song copying and the number of tutor song bouts.

### Song preference predicted song learning

We next tested whether the degree to which the pupils *preferred* control song over OTA song predicted the degree to which they *learned* control song better than OTA song. We found a highly significant, positive correlation between the average preference for control song and the learning difference score, calculated as the similarity of the pupil’s song to control song minus its similarity to OTA song (*Rho* = 0.943; *p* = 0.005; Fig. 4C). Preference for control song was also significantly positively correlated with learning of control song (*Rho* = 0.829, *p* = 0.042) and significantly negatively correlated with learning of OTA song (*Rho* = -0.886, *p* = 0.019). When we ran these correlations using preference data within each learning phase, we found that the learning difference score was predicted by preference significantly only during the auditory learning phase (*Rho* = 0.829, *p* = 0.042) but there were trends during the plastic song phase and the crystallization phase (both phases *Rho* = 0.771; *p* = 0.072).

## Discussion

Here, we have shown that OTR antagonism during song tutoring both reduced preferences for that tutor’s song and inhibited learning of that song. This result is consistent with our hypothesis that OTR stimulation increases attraction to and attention to tutors during an early critical period, facilitating song learning. These findings add to a small but growing literature suggesting that OTR may promote early social bonds with caregivers, thus creating conditions conducive to enhanced social learning. Such a mechanism may be particularly relevant in the context of human language learning, which requires attention to a targeted social object such as a caregiver’s face or voice. In this way, vocal development may be ‘gated’ by dynamic social interactions (Baran, 2017; Kuhl, 2007; Syal & Finlay, 2011; Woolley & Moore, 2011).

In addition to its effects on preferences and learning, OTA also measurably altered behavior during tutoring itself. We found that when treated with OTA, the pupils were less likely to engage in behaviors that may indicate interest in the tutor, namely approaching the tutor and pecking on the cage wall. We also found that OTA inhibited preening, a behavior done while sitting quietly. Both preening and quiet stillness were previously shown to predict the accuracy of song copying in zebra finches, perhaps because these behaviors facilitate listening (Chen et al., 2016; Houx et al., 2000).

One of our most interesting findings was that treatment of the *pupils* affected the behavior of *tutors*. When presented with OTA-treated pupils, tutors sang at lower rates than when presented with control-treated pupils. Other researchers have hypothesized that tutor song rate may be affected by pupil behavior; however there is little information on what those behaviors might be. In an exhaustive study of contingent tutor-pupil behaviors, Houx & ten Cate (1998) found no evidence that tutor song was predicted by any overt behavior from the pupil. Because of our small sample size and because we scored only a limited number of behaviors, we could not isolate a particular behavior in pupils that predicted tutor song rate. Nonetheless, our results suggest that tutors may attend specifically to oxytocin-dependent behaviors of their pupils.

In another recent study using the same operant conditioning assay, we quantified preference for the male caregiver’s (father’s) song in juvenile male zebra finches (Rodriguez-Saltos et al., 2021). In that study, pupils were naturally reared by their parents, both mother and father. At 35 dph, these pupils were transferred into an operant chamber with two keys, one of which was associated with the father’s song and the other with the song of a familiar “neighbor” that they could hear but not interact with in their home cage during rearing. In that study, we found that the degree to which the pupils preferred the song they ultimately sang (which in that study was always the father’s song) strongly predicted the quality of learning of that song. That finding was replicated here; the degree to which each pupil preferred the control song over the OTA song significantly predicted the degree to which the pupil learned the control song better than the OTA song (Fig. 4C). Even though our sample size was small, this effect was dramatic. Together, these findings suggest that the ‘incentive salience’ of a signal early during development—in other words, its attractiveness—facilitates social learning of that signal.

Our current findings recapitulate the findings of Rodriguez-Saltos et al. (2021) in several other ways, including the shape of the developmental trajectory of song preference. In that study, the pupils demonstrated a significant preference for the father’s song during the auditory phase of learning. Then, at around 55 dph, their preferences shifted dramatically to the neighbor’s song, reaching statistical significance at 57-65 dph. Then, these males showed no preference for either song until crystallization, when there was another significant peak in preference for the neighbor’s song. Thus, the trajectory of preference for the father’s song, which is what all of the pupils ultimately sang in that study, was nearly identical to the trajectory of preference for control song in the current study. In other words, the pupils in the current study appeared to treat the control song as if it were the song of their father, and the OTA song as if it were the song of a familiar male with whom they had no social bond.

The timing of the shift in preference from the father’s song to the neighbor’s song, or in this study, from the control song to the OTA song, is important. In both studies, the preference for the song that was ultimately learned crossed the 0.5 mark, in other words shifted to the other song, precisely at the time that juveniles leave the auditory learning phase and begin singing plastic song (Fig. 1). This time, around 55 dph, is a time of major change in the life of a juvenile zebra finch, potentially corresponding to puberty (Ball & Wade, 2013; Zann, 1996). The young male finch is unlikely to memorize a new song that he hears after this time, preferring to practice the song that he has already chosen to learn (Eales 1985; Gobes et al., 2019). His song rate increases dramatically (Johnston et al., 2002) and the syllables in his song begin to resemble those in tutor song (Kollmorgen et al., 2020). At the same time, he begins social exploration, spending more time away from the parents and instead interacting with unrelated finches (Adkins-Regan & Leung, 2006). Thus, we interpret the shift in preference as a sign of social exploration, or wanting to hear songs other than the one the pupil has himself chosen to learn.

Post-hatch day 55 is significant also because at this time, OTR expression in the brain undergoes dramatic changes. In a previous study (Davis et al., 2019), we quantified the expression of OTR mRNA in four regions relevant to sociality or song learning: the lateral septum, the auditory forebrain, HVC, and the dorsal arcopallium, which makes up part of the ‘shelf’ region around another song nucleus, the robust nucleus of the arcopallium. We found that in every region we looked at, OTR mRNA expression declined precipitously at 55 dph. Expression then rebounded to previous levels by 65 or 75 dph. We hypothesize that this striking decrease, which was widespread in the brain yet limited to a relatively short time window, plays a role in the co-occurring shift in social preferences and behaviors.

This is not the first study to show effects of nonapeptides on song learning. Baran et al. (2016; 2017) treated zebra finch hatchlings with daily intracranial injections of Manning Compound, a V1aR antagonist, and tracked their behavior until maturity. Treated birds exhibited atypical social development and showed no clear preference to affiliate with their parents vs. unfamiliar conspecifics. Treated males were also impaired in their song acquisition, and the degree of impairment was correlated with measures of atypical social behaviors (Baran et al., 2017). Therefore, V1aR may play a role in early attachment and song learning. The binding pattern of the antagonist used in this study, ornithine vasotocin, resembles the distribution of OTR mRNA, not V1aR mRNA, regardless of age (Davis et al., 2019; Leung et al., 2011). We therefore believe that if the antagonist used in this study is crossing into the brain, it is binding primarily to OTR rather than V1aR or another type of nonapeptide receptors. We cannot, however, make predictions about the extent to which binding of the two endogenous ligands, mesotocin and vasotocin, is affected by this antagonist. Both nonapeptides are predicted to bind OTR (Leung et al., 2011).

One limitation of this study was that we administered OTA peripherally rather than directly into the brain. The antagonist we used has been shown by others to affect social behavior in zebra finches when injected peripherally (Goodson et al., 2009; Pedersen & Tomaszycki, 2012); however it will be necessary to conduct central infusions, targeting particular receptor populations, to better understand the role of OTR in social reward during song learning.

Another potential limitation is that, because we were primarily interested in the effects of oxytocin antagonism on tutor choice, we included only males in this study. Others have shown that females also show preferences for their father’s song in choice tests. In adulthood, these preferences are just as strong as in males (Riebel et al., 2002) or even stronger (Fujii et al., 2021). OTR expression, namely in song control regions, is higher in males than females during the auditory learning phase (Davis et al., 2019), suggesting that some OTR populations could be involved in song learning specifically. Future experiments should explore the possibility that social reward, particularly attraction to song, could rely on different OTR populations in females and males.

## Materials and Methods

### Animals

All procedures were approved by the Emory University Institutional Animal Care and Use Committee. Juveniles used in the study (n = 9), or “pupils”, were produced by a breeding colony at Emory University and hatched in 2018-2019. All birds were housed under a 12-hour-light/12-hour dark cycle. Cages (14 × 14 × 16 inches) were kept individually in sound-attenuating chambers (interior dimensions 22 x16 × 23 inches, Colbourn Instruments, H10-24TA) to prevent exposure to adult songs. Birds were given water, seed, cuttlebone, and greens *ad libitum*. To discourage nestlings from developing a side bias that could affect our ability to measure song preference in the operant assay, all food cups and water bottles were matched by color and positioned symmetrically in the cage. The father of the nestlings was removed at four dph, counting from the hatch date of the oldest nestling, leaving the female to provide all parental care. After the father’s removal, therefore, hatchlings did not hear song until tutoring began (Fig. 1). To ensure that all the pupils we selected were male and, therefore, could learn to sing, the sex of nestlings was determined using PCR analysis of DNA from blood samples (Griffiths, et al., 1998). Sex was later confirmed by plumage.

### Song tutoring

We selected adult male tutors with high song rates. Song rate was determined by placing the cage of an adult male next to the cage of either an adult female or a juvenile male for one hour and noting the rate of singing. To limit potential bias towards learning one of the two tutor songs, each pupil was matched to tutors with songs dissimilar to that of the pupil’s genetic father. This dissimilarity was assessed by visually inspecting spectrographs created in Audacity. We looked for shared “syllables”, or short, stereotyped sounds that are separated by a brief period of silence (Zann, 1996). We matched each pupil to tutors that shared no more than two syllables with the pupil’s father (Range 0-2, M = 0.667, SD = 0.623). Moreover, for each pupil, the two tutors shared no more than two syllables with each other (Range, 0-2, M = 0.6, SD = 0.8). The shared syllables were primarily stereotyped, call-like sounds that typically vary little among individuals (Zann, 1996).

Tutoring sessions took place at 27, 29, 31, and 33 dph, which corresponds to a period of high receptivity to tutoring (Böhner, 1990). Thirty minutes before each of four tutoring trials, pupils were treated with either saline (control) or an oxytocin receptor antagonist [d(CH2)51,Tyr(Me)2,Thr4,Orn8,des-Gly-NH29] -Vasotocin trifluoroacetate salt (Bachem), dissolved in saline. This antagonist was previously shown to reduce social approach (Goodson et al., 2009) and pair-bond formation (Pedersen & Tomaszycki, 2012) when administered peripherally in zebra finches. In this study, both OTA and saline were administered via a 0.05 mL subcutaneous injection into the shoulder area between the feather tracts. Trials were separated by ∼48h and the treatments, OTA or saline, were alternated. The order in which the pupils received each treatment was counterbalanced, such that some pupils received OTA first and some received saline first.

For each pupil, each of the two treatments was always associated with the same tutor; in other words, there was an “OTA tutor” and a “control tutor” for each pupil. Tutor pairs were each used for two pupils, with the identity of the control tutor and OTA tutor reversed. Pupils spent two tutoring sessions with each of their tutors, for a total of four sessions per pupil. Each tutoring session took place inside a walk-in sound-attenuating booth and lasted about an hour (London & Clayton, 2008). Immediately after receiving the subcutaneous injection of OTA or saline, a pupil was placed into a testing cage, brought into a booth, and given 30 minutes alone to acclimate to this environment before the tutoring session commenced. The tutor was given about 15 minutes to habituate in a separate, identical booth before being moved, in its cage, to the booth with the pupil. The tutor and pupil remained in separate cages during the tutoring sessions but could interact visually and vocally. We defined the start of the tutoring session as the moment when the experimenter left the room after placing the tutor in the booth with the pupil. Audio and video recordings were made of all tutoring sessions using a camera placed inside the booth.

We took steps to balance the number of times each pupil heard each tutor’s song during tutoring sessions, such that each pupil heard roughly the same number of songs from each of the two tutors. On the basis of our calculations of each tutor’s singing rate, which was made when selecting the tutors, we estimated the approximate number of songs that we expected each pupil to hear during each of the four sessions. If this number was not reached by 45 minutes into the session, recordings of that tutor’s song were played from a speaker at 75 dB during the last ∼15 minutes of the tutoring session. In instances when the tutor sang at a rate higher than anticipated, tutoring sessions were cut a few minutes short in order to control the number of songs heard by the pupil. The average session duration was 58.6 min ± 5.0 min. After each session, the pupil and the tutor were brought back to their respective home cages.

### Behavioral observations

After all the trials were completed, the recordings were observed using Behavior Observation Research Interactive Software (BORIS, v.7.9.19). We focused on behaviors that others have previously found to be predictive of song learning, such as approaching the tutor (Houx et al., 2000) and paying attention to the tutor (Chen et al., 2016). To quantify approach to the tutor, we scored two behaviors. First, we quantified the percentage of total time during the session that the pupil spent close to the tutor, defined as within an area ≤12 cm from the wall of its own cage that faced the tutor’s cage. This area of the cage was designated as the “tutor zone.” Our second measure of approach was “pecks to tutor”, which was counted each time the pupil approached the wall of its own cage closest to the tutor and pecked at that wall.

Attention during tutoring, or more specifically, lack thereof, has been operationalized by others as the level of intense activity, such as flying, and number of vocalizations by the pupil, which both negatively predict learning (Chen et al., 2016). To quantify flying, we scored a behavior that occurred frequently in the most active pupils: two-foot wall contacts, in other words when the pupil flew and landed with both of its feet on any side wall of the cage. In addition, we counted all vocalizations made by the pupils. These vocalizations did not include song, as the tutoring sessions took place largely before juveniles begin producing even immature song (Immelmann, 1969; Fig. 1). As a final measure of attention, we also scored preening, a grooming behavior that we defined as when the pupil used its beak to clean its feathers or another part of its body. Preening during tutoring has been found to be positively associated with learning, possibly because the bird sits quietly while doing it (Houx et al., 2000).

In addition to the pupil’s behavior, we also scored singing by the tutor. We defined a tutor song bout as an instance of continuous singing with at least a one-second gap between bouts. Because tutor songs vary slightly in duration and we were more interested in the number of times the pupil heard the song than the total duration of singing, we scored a song bout as a point event.

To test for effects of OTA treatment on behavior during tutoring, we used linear mixed models (LMMs) in R (v4.04). Separate tests were conducted for each measure. In all models, pupil ID was included as a random effect to accommodate the within-subjects design. For all variables, we used the following model: ∼ Trial + Treatment + (1 | Pupil ID), which allowed us to control for the effects of Trial (i.e. the first or second tutoring session for each treatment). To perform the LMM analyses, we used the *lmer* function in the lme4 package (Bates et al., 2014). To test for a significant effect of Treatment on each variable of interest, we performed a likelihood ratio test using the *anova* function to compare the full model to a reduced model with Treatment removed. As a measure of effect size, we calculated Cohen’s d_av_ as described by Cumming (2012; see also Lakens, 2013), which takes into account the within-subjects design.

### Auditory stimuli

A unit of song contains two consecutive repeats of a highly stereotyped motif (Woolley & Moore, 2011; Zann, 1996). To create the playbacks used during tutoring sessions (see *Song Tutoring*, above) and in the preference assay (see *Operant Assay*, below), two consecutive motifs from each tutor were isolated in audio files. Male zebra finches typically produce a series of call-like introductory notes immediately before beginning to sing; these introductory notes were excluded. The files were then cleaned using Audacity as described by Rodriguez-Saltos et al. (2021).

### Operant assay of song preference

Keys for the operant assay were assembled from microswitches (ZF Electronics, #D429-R1ML-G2). Pupils were removed from their home cage at 35 dph, 48 hours after the last tutoring session concluded, and housed singly in a cage inside a sound-attenuating chamber. By this age, juvenile zebra finches are able to feed themselves independently (Zann, 1996). The operant set-up was as described by Rodriguez-Saltos et al. (2021). Two keys, accessible from a perch, were placed on opposing walls. (Fig. 3A). Each key was paired with one of two speakers, which were located inside the chamber on opposing sides. The speakers were both set to play song at 75 dB as measured at the center of the cage. A small mirror was affixed to the center of the cage’s back wall to provide enrichment.

The operant assay ran on a probabilistic schedule of reinforcement controlled by the custom-written software program SingSparrow (Rodriguez-Saltos et al., 2021). The program allowed the pupil to “self-tutor” by pressing a key to trigger a playback. When a playback was triggered, one song composed of two motifs played through the associated speaker. Juveniles have been shown to learn song effectively using this self-tutoring method, even with hearing as few as 15 songs each day (Tchernichovski et al., 1999). Our operant schedule was designed to balance exposure to the two tutor songs, limiting each to 30 playbacks, while still allowing us to quantify preference for a particular song each day (Fig. 3B). This “self-balancing” schedule was designed as follows: both keys could play both songs, but each key was associated with a higher probability of playing one song or the other. As a key was pressed repeatedly, the probability that it would play its associated song decreased stepwise from 100 percent to 50 percent. Each key was, however, always more likely to play its associated song than was the other key, as long as the quota of 30 for that song had not been met. Additionally, whenever the pupil switched between keys, the first press following the switch always played that key’s associated song. After 30 playbacks had been triggered of one song, both keys played only the other song – the song in deficit - until all 60 songs were exhausted. At this point, the pupil could continue to press the keys, but no song would play.

On 36 dph, or the day after the pupil was isolated in an operant chamber, the keys were activated to begin the operant assay at lights-on. The assay ran through 99 dph, with the keys being reactivated every day at lights-on. The number of presses on each key was automatically logged between lights-on and lights-off each day, as well as which song played. To detect possible side biases, reversals were run as described by Rodriguez-Saltos et al. (2021).

### Analysis of song preference

The strength of a pupil’s preference for a particular song was determined by counting the number of presses per day on each key (Rodriguez-Saltos et al., 2021). Key presses were removed if they were triggered accidentally by the caretaker or by the bird while care was being provided. Data were excluded from any days during which a pupil’s total number of presses was fewer than five; these days happened early during training when the pupils were still learning the task. Daily counts of key presses were then filtered to include only those presses that occurred up to the point in the schedule when the quota of 30 playbacks for one song had been reached. After this point, the pupil no longer had a choice of which song to hear, making key presses less behaviorally meaningful.

After the data were cleaned as described above, we calculated daily “preference scores” for each pupil as a proportion of presses on one of the keys, e.g.: Key A presses / (Key A + Key B presses). Preference scores were calculated as the preference for the key associated with the song of the control tutor (“control song”; preference for OTA song is preference for control song subtracted from 1). To reconstruct the average trajectory of preference for control song, we fitted a generalized additive model to our data (Wood, 2011). The dependent variable was preference for the control song and the independent variable was age in days. Pupil identity was modelled as a random effect. To constrain predicted values to the interval [0,1], we modelled the dependent variable using a beta regression link. The relationship between preference and age was modelled using a thin-plate regression spline (Wood, 2003). The model was fitted using the library *mgcv* (Wood, 2017) in R (R Core Team, 2019). *Mgcv* produced a 95% confidence interval of the trajectory by multiplying the standard error of the trajectory by two, subtracting this result from the trajectory to find the lower bound, and summing it to the trajectory for upper bound (Wood, 2017). At any given age the pupils significantly preferred the control song over OTA song if the trajectory was above 0.5 and the confidence interval excluded that value.

### Quantification of song learning

When the pupils reached adulthood, their songs were recorded and cleaned as described above. Songs were recorded at 101-103 dph, the point at which pupils have completed their vocal development and produced a crystallized song that does not change much thereafter (Johnson et al., 2002; Price, 1979; Tchernichovski et al., 2001; Zann, 1996). Song recordings were imported as .WAV files into Sound Analysis Pro 2011 (SAP). SAP generates sound spectrograms allowing visualization of a song’s acoustic features such as its amplitude, entropy, and pitch (Tchernichovski et al., 2000). To calculate similarity between tutor and pupil songs, we segmented songs into syllables on the basis of an amplitude threshold (Rodriguez-Saltos et al., 2021; Tchernichovski et al., 2000). Thresholds were adjusted for every song because each one was sung at a different distance from the microphone; thus, song-specific amplitude thresholds allowed for each song to be properly segmented. Similarity scores were calculated using SAP’s “Song Similarity” tool, which compares the acoustic features of one song’s syllables to another and quantifies similarity (Tchernichovski et al., 2000). Similarity scores were calculated for each pupil-tutor pair using three to six song exemplars per pupil. We used the maximum similarity score for each pupil-tutor pairing for the analysis (Shi et al., 2017; 2018). To test for an effect of OTA treatment on song learning, we used LMMs as described above (*Behavioral Observations*) using the following model: ∼ Treatment + (1|Pupil ID), where Pupil ID was included as a random effect.

To test for relationships among factors such as song preferences, song learning, and the number of live tutor songs heard, we used Spearman correlation tests.

## Acknowledgments

We are indebted to Gordon Ramsay and Ami Klin, who listened to our ideas and encouraged us to develop the zebra finch as a new model of social reward. We are grateful to Jocelyne Bachevalier, Patricia Bauer, Aubrey Kelly, Jennifer Merritt, and Kristin Wendland for feedback on previous versions of this manuscript, as well as to Isabel Fraccaroli and Erik Iverson for technical assistance. We also thank Erich Jarvis and Laura Carruth for providing founders for our zebra finch colony. This work was supported by NIH 1R21MH105811-01A1 to DLM and by the Silvio O. Conte Center for Oxytocin and Social Cognition, 2P50MH100023. The authors declare no conflicts of interest.

## Notes

### Competing Interest Statement

The authors have declared no competing interest.

## References

Adkins-Regan, E. & Leung, C. H. (2006). Sex steroids modulate changes in social and sexual preference during juvenile development in zebra finches. Hormones and Behavior,

Ball, G. F. & Wade, J. (2013). The value of comparative approaches to our understanding of puberty as illustrated by investigations in birds and reptiles. Hormones and behavior, 64(2), 211–214.

Baran, N. M. (2017). Sensitive periods, vasotocin-family peptides, and the evolution and development of social behavior. Frontiers in Endocrinology, 8, 189.

Baran, N. M., Peck, S. C., Kim, T. H., Goldstein, M. H., & Adkins-Regan, E. (2017). Early life manipulations of vasopressin-family peptides alter vocal learning. Proceedings of the Royal Society B: Biological Sciences, 284(1859), 20171114.

Baran, N. M., Sklar, N. C., & Adkins-Regan, E. (2016). Developmental effects of vasotocin and nonapeptide receptors on early social attachment and affiliative behavior in the zebra finch. Hormones and Behavior, 78, 20–31.

Bates, D., Mächler, M., Bolker, B., & Walker, S. (2014). Fitting linear mixed-effects models using lme4. Cornell University, arXiv: 1406.5823.

Böhner, J. (1990). Early acquisition of song in the zebra finch, Taeniopygia guttata. Animal Behaviour 39, 369–374.

Chen, Y., Matheson, L. E., & Sakata, J. T. (2016). Mechanisms underlying the social enhancement of vocal learning in songbirds. Proceedings of the National Academy of Sciences, 113, 6641–6646.

Cousineau, D. (2005). Confidence intervals in within-subject designs: A simpler solution to Loftus and Masson’s method. Tutorials in Quantitative Methods for Psychology, 1(1), 42–45.

Cumming, G. (2012). Understanding the new statistics: Effect sizes, confidence intervals, and meta-analysis. New York, NY: Routledge.

Davis, M. T., Grogan, K. E., & Maney, D. L. (2019). Expression of oxytocin receptors in the zebra finch brain during vocal development. bioRxiv, 739623.

Deshpande, M., Pirlepesov, F., & Lints, T. (2014). Rapid encoding of an internal model for imitative learning. Proceedings of the Royal Society of London B: Biological Sciences, 281(1781), 20132630.

Eales, L. A. (1987). Do zebra finch males that have been raised by another species still tend to select a conspecific song tutor? Animal Behaviour, 35(35), 1347–1355.

Eales, L. A. (1985). Song learning in zebra finches: Some effects of song model availability on what is learnt and when. Animal Behaviour, 33(4), 1293–1300.

Fujii, T. G., Ikebuchi, M., & Okanoya, K. (2021). Sex differences in the development and expression of a preference for familiar vocal signals in songbirds. PLoS ONE, 16(1), e0243811.

Gobes, S. M. H., Jennings, R. B., & Maeda, R. K. (2019). The sensitive period for auditory-vocal learning in the zebra finch: Consequences of limited-model availability and multiple-tutor paradigms on song imitation. Behavioural Processes, 163, 5–12.

Goodson, J., Schrock, S., Klatt, J., Kabelik, D., & Kingsbury, M. (2009). Mesotocin and nonapeptide receptors promote estrildid flocking behavior. Science, 325, 862–865.

Gordon I, Martin C, Feldman R, & Leckman JF (2011). Oxytocin and social motivation. Developmental Cognitive Neuroscience 1, 471–493.

Griffiths, R., Double, M. C., Orr, K., & Dawson, R. J. G. (1998). A DNA test to sex most birds. Molecular Ecology, 7, 1071–1075.

Gubrij, K. I., Chaturvedi, C. M., Nawab, A., Cornett, L. E., Kirby, J. D., Wilkerson, J., … & Baeyens, D. A. (2005). Molecular cloning of an oxytocin-like receptor expressed in the chicken shell gland. Comparative Biochemistry and Physiology, 142(1), 37–45.

Hammock EA (2014). Developmental perspectives on oxytocin and vasopressin. Neuropsychopharmacology, 40, 24–42.

Houx, B. & ten Cate, C. (1998). Do contingencies with tutor behaviour influence song learning in zebra finches? Behaviour, 135(135), 599–614.

Houx, B., ten Cate, C., & Feuth, E. (2000). Variations in zebra finch song copying: An examination of the relationship with tutor song quality and pupil behaviour. Behavioural Biology, 137, 1377–1389.

Immelmann, K. (1969). Song development in the zebra finch and other estrildid finches. In R. A. Hinde (Ed.), Bird Vocalisations (pp. 61–74). Cambridge: Cambridge University Press.

Johnson, F., Soderstrom, K., & Whitney, O. (2002). Quantifying song bout production during zebra finch sensory-motor learning suggests a sensitive period for vocal practice. Behavioural Brain Research, 131(1), 57–65.

Kelly, A. M., & Goodson, J. L. (2014). Social functions of individual vasopressin–oxytocin cell groups in vertebrates: What do we really know?. Frontiers in Neuroendocrinology, 35(4), 512–529.

Kollmorgen, S., Hahnloser, R. H. R., & Mante, V. (2020). Nearest neighbours reveal fast and slow components of motor learning. Nature, 577(7791), 526–530.

Kuhl, P. K. (2007). Is speech learning ‘gated’ by the social brain? Developmental Science, 10(10), 110–120.

Lakens, D. (2013). Calculating and reporting effect sizes to facilitate cumulative science: A practical primer for t-tests and ANOVAs. Frontiers in Psychology, 4, 863.

Leung, C. H., Abebe, D. F., Earp, S. E., Goode, C. T., Grozhik, A. V., Mididoddi, P., & Maney, D. L. (2011). Neural distribution of vasotocin receptor mRNA in two species of songbird. Endocrinology, 152(12), 4865–4881.

Leung, C.H., Goode, C.T., Young, L.J., & Maney, D.L. (2009). Neural distribution of nonapeptide binding sites in two species of songbird. Journal of Comparative Neurology, 513,197–208.

London, S. E. & Clayton, D. F. (2008). Functional identification of sensory mechanisms required for developmental song learning. Nature Neuroscience, 11(5), 579–586.

Loveland, J.L., Stewart, M.G., Vallortigara, G. (2019). Effects of oxytocin-family peptides and substance P on locomotor activity and filial preferences in visually naïve chicks. European Journal of Neuroscience, 50, 3674–3687.

Maney, D. L. & Rodriguez-Saltos, C. A. (2016). Hormones and the incentive salience of bird song. In Hearing and Hormones (Vol. 57, pp. 101–132). New York: Springer International Publishing.

Morey, D. R. (2008). Confidence intervals from normalized data: A correction to Cousineau (2005). Tutorials in Quantitative Methods for Psychology, 4(2), 61–64.

Pedersen, A. & Tomaszycki, M. L. (2012). Oxytocin antagonist treatments alter the formation of pair relationships in zebra finches of both sexes. Hormones and Behavior, 62(2), 113–119.

Price, P.H. (1979). Developmental determinants of structure in zebra finch song. Journal of Comparative and Physiological Psychology, 93(2), 260–277.

Riebel, K., Smallegange, I. M., Terpstra, N. J., & Bolhuis, J. J. (2002). Sexual equality in zebra finch song preference: evidence for a dissociation between song recognition and production learning. Proceedings of the Royal Society of London. Series B: Biological Sciences, 269(1492), 729–733.

Rodriguez-Saltos, C. A., Bhise, A., Karur, P., Khan, R. N., Lee, S., Ramsay, G., & Maney, D. L. (2021). Song preferences predict the quality of vocal learning in zebra finches. bioRxiv, 2021.06.01.446570.

Ross, H.E. & Young, L.J. (2009). Oxytocin and the neural mechanisms regulating social cognition and affiliative behavior. Frontiers in Neuroendocrinology, 30, 534–547.

Shi, Z., Piccus, Z., Zhang, X., Yang, H., Jarrell, H., Ding, Y., … & Li, X. (2018). MiR-9 regulates basal ganglia-dependent developmental vocal learning and adult vocal performance in songbirds. eLife, 7, e29087.

Shi, Z., Tchernichovski, O., & Li, X. (2017). Studying the mechanisms of developmental vocal learning and adult vocal performance in zebra finches through lentiviral injection. Bio Protocol, 8(17).

Syal, S. & Finlay, B. L. (2011). Thinking outside the cortex: Social motivation in the evolution and development of language. Developmental Science, 14(2), 417–430.

Tchernichovski, O., Mitra, P. P., Lints, T., & Nottebohm, F. (2001). Dynamics of the vocal imitation process: How a zebra finch learns its song. Science, 291(5513), 2564–2569.

Tchernichovski, O., Nottebohm, F., Ho, C. E., Pesaran, B., & Mitra, P. P. (2000). A procedure for an automated measurement of song similarity. Animal Behaviour, 59(6), 1167–1176.

Tchernichovski, O., Lints, T., Mitra, P. P., & Nottebohm, F. (1999). Vocal imitation in zebra finches is inversely related to model abundance. Proceedings of the National Academy of Sciences, 96(22), 12901–12904.

ten Cate, C. (1986). Listening behaviour and song learning in zebra finches. Animal Behaviour, 34(4), 1267–1268.

Theofanopoulou, C., Gedman, G., Cahill, J. A., Boeckx, C., & Jarvis, E. D. (2021). Universal nomenclature for oxytocin–vasotocin ligand and receptor families. Nature, 592(7856), 747–755.

Theofanopoulou, C., Boeckx, C., & Jarvis, E. D. (2017). A hypothesis on a role of oxytocin in the social mechanisms of speech and vocal learning. Proceedings of the Royal Society of London B: Biological Sciences, 284, 20170988.

Vaidyanathan, R. & Hammock, E.A. (2017). Oxytocin receptor dynamics in the brain across development and species. Developmental Neurobiology, 77, 143–157.

Wood, S. (2017). Generalized Additive Models: An Introduction with R, Second Edition. CRC Press. https://www.routledge.com/Generalized-Additive-Models-An-Introduction-with-R-Second-Edition/Wood/p/book/9781498728331.

Wood, S. N. (2011). Fast stable restricted maximum likelihood and marginal likelihood estimation of semiparametric generalized linear models. Journal of the Royal Statistical Society: Series B (Statistical Methodology), 73(1), 3–36.

Wood, S. N. (2003). Thin plate regression splines. Journal of the Royal Statistical Society: Series B (Statistical Methodology), 65(1), 95–114.

Woolley, S. M. N. & Moore, J. M. (2011). Coevolution in communication senders and receivers: vocal behavior and auditory processing in multiple songbird species. Annals of the NY Academy of Sciences, 1225, 155–165.

Zann, R. A. (1996). The zebra finch: A synthesis of field and laboratory studies. Oxford University Press.

