## Supplemental Figures for "Oxytocin receptor antagonism during song tutoring in zebra finches reduces preference for and learning of the tutor’s song"

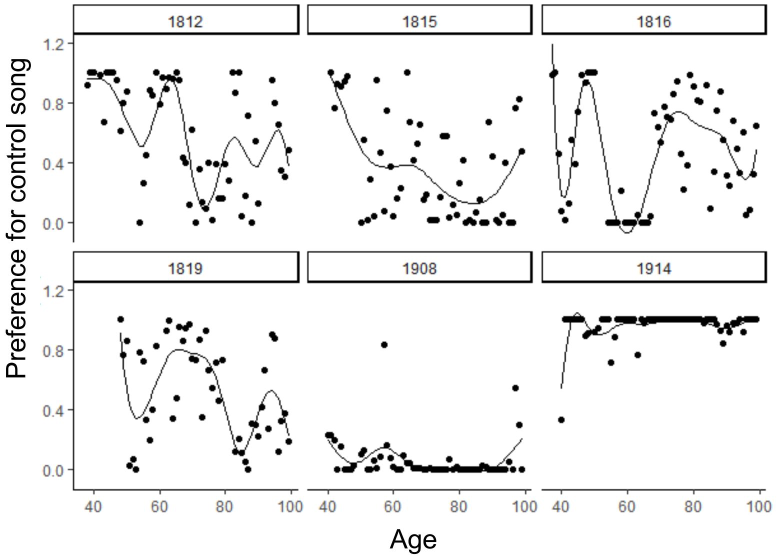


**Fig. S1. Individual trajectories of song preference.** The developmental trajectories of preference are shown for the six pupils in this study that completed the key-pressing assay. The preference for control song was calculated as the proportion of presses for the key associated with that song. The dots in each plot are the daily preference scores for each bird. A smooth trajectory was calculated by applying LOESS to the datapoints. Despite the original data consisting of proportions, and thus bound to the interval [0-1], LOESS may fit values slightly outside that interval.

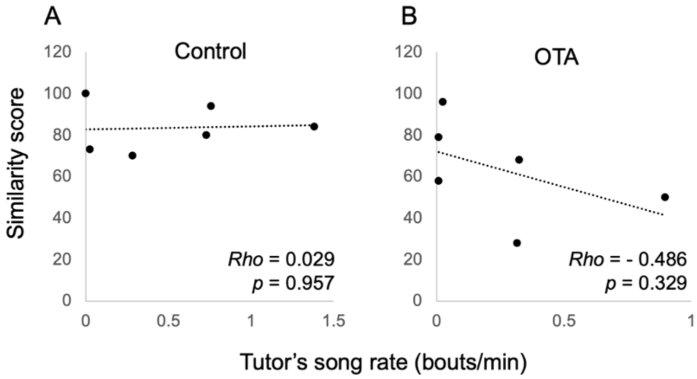


**Fig. S2. Lack of correlations between tutor song rate and learning.** The tutor's song rate did not predict the degree to which that song was learned, either for the control tutor (A) or the OTA tutor (B).
